# Increased hippocampal GABAergic inhibition after long-term high-intensity sound exposure

**DOI:** 10.1101/507004

**Authors:** AOS Cunha, JL de Deus, CC Ceballos, RM Leão

**Affiliations:** Department of Physiology, FMRP, University of São Paulo, Ribeirão Preto, SP. Brazil

**Keywords:** hippocampus, high intensity sound, GABA, glutamate, neurotransmission

## Abstract

Exposure to loud sounds has been related to deleterious mental and systemic effects in addition to auditory maladies. Hippocampal function has been shown to be affected to either high intensity sound exposure or long-term sound deprivation. Hippocampal long-term potentiation (LTP) is inhibited after 10 days of daily exposure to 2 minutes of high-intensity noise (110 dB), in the hippocampi of Wistar rats. He we investigate how the glutamatergic and GABAergic neurotransmission mediated by ionotropic receptors is affected by the same protocol of high intensity sound exposure. We found that while the glutamatergic transmission both by AMPA/kainite and NMDA receptors in the Schaffer-CA1 synapses is largely unaffected by long-term exposure to high intensity sound, the amplitude of the inhibitory GABAergic currents is potentiated, but not the frequency of the both spontaneous and miniature currents. We conclude that GABAergic transmission is potentiated at the post-synaptic level in the hippocampal CA1 pyramidal neurons after a prolonged exposure to short periods of high-intensity sound. This effect could be an important factor for the reduced LTP in the hippocampi of these animals after high intensity sound exposure, and demonstrated that prolonged exposure to high- intensity sound can affect hippocampal inhibitory transmission and consequently its function.

## Introduction

Exposure to loud noises has been related to several deleterious mental and systemic effects [1,2], in addition to auditory maladies as deafness, hyperacusis and tinnitus [3,4]. Exposure to loud sound is an increasing source of acoustic pollution, both from occupational and recreational sources, and health problems related to sound exposure are increasingly common, even in juveniles [5,6,7]. However while the effects of high-intensity sound exposure in the central auditory neurons have been extensively studied, the effects in other brain areas, especially in areas related to cognition and emotions, are less known.

The hippocampus is a region traditionally implicated in the formation of declarative and spatial memories and is connected to the auditory system indirectly from the frontomedial cortex, insula and amygdala [8], and recently a direct contact between the auditory cortex and CA1 region was identified [9]. This pathway is implicated in the formation of long-term auditory memories [10], and auditory cues can be used in the formation of spatial memories [11]. Hippocampal place cells can be activated by a task using auditory dimension cues [12] and recently it was demonstrated that sound stimulation evoke excitatory and inhibitory currents in CA3 pyramidal neurons *in vivo* [13].

Accordingly, sound stimulation or sound deprivation affect the hippocampus. Prolonged (2 hours/ day for 3-6 weeks) moderate (80 dB) sound exposure, impairs spatial memory in mice and increases oxidative damage and tau phosphorylation in the hippocampus [14,15]. Acute traumatic noise (106-115 dB, 30-60 minutes) alters place cell activity in the hippocampus [16] and increases *arc* expression, an immediate early gene related to synaptic plasticity, in the hippocampi of rats [17]. Work from our group has shown that long-term (10 days, 2 minute a day) stimulation with high intensity broadband sound inhibits LTP in the Schaffer-CA1 hippocampus of rats [18] and hyperpolarizes CA1 pyramidal neurons by decreasing the expression of the h current (I_h_), while increasing the firing of these neurons by an unknown ionic mechanism [19]. The effect on LTP, but not on the membrane intrinsic properties of CA1 neurons, was also observed after a single one-minute 110 dB broadband sound stimulus [20]. In order to investigate the possible mechanisms underlying the inhibition of LTP by high intensity sound, we investigated using whole-cell patch-clamp recordings *in vitro*, the excitatory and inhibitory neurotransmission in CA1 pyramidal neurons from animals subjected to long-term exposure to episodes of high-intensity broadband noise.

## Material and Methods

### Animals

All experimental procedures involving animals were designed according to rules for animal research from the National Council for Animal Experimentation Control (CONCEA) and approved Commission for Ethics in Animal Experimentation (CEUA) at the University of São Paulo at Ribeirão Preto (protocols 015/2013 and 006/2-2015). Male Wistar rats (60-80 days) were obtained from the central animal facility of the University of São Paulo-Ribeirão Preto Campus and kept in the rat animal facility of the School of Medicine of Ribeirão Preto until the day of use. The animals were kept in Plexiglas cages (2-3 animals per cage), food and water ad libitum and 12-h dark/light cycle (lights on at 7:00 a. m.) and controlled temperature (22 °C).

### Sound stimulation protocol

Our protocol was previously described in [18]. Briefly, animals were placed in an acrylic arena (height: 32 cm, diameter: 30 cm) located inside an acoustically isolated chamber (45 x 45 x 40 cm, 55 dB ambient noise) where, after one minute of habituation, they were submitted to a one-minute episode of 110-dB sound stimulus (a digitally modified recording of a doorbell, spanning frequencies from 3 to 15 kHz) [21]. The animals were kept in the stimulation chamber for one more minute, and then returned to their home cages. In case the animals presented seizures [18] they were discarded from the study. This protocol was repeated for 10 days, twice a day (8-9 am and 4-5 pm). The animals rested for 10-14 days after the last session before sacrifice. The control group was placed in the box for the same amount of time and not subjected to sound stimulation.

### Hippocampal slices

Animals were anesthetized with isoflurane, decapitated and the brains rapidly removed and placed in an ice-cold solution containing (mM): 87 NaCl, 2.5 KCl, 25 NaHCO_3_, 1.25 NaH_2_PO_4_, 75 Sucrose, 25 Glucose, 0.2 CaCl_2_, 7 MgCl2, bubbled with 95% O2 and 5% CO_2_. The brain hemispheres were separated and glued with cyanoacrylate glue to a support and placed inside the cutting chamber of a vibratome (1000 plus, Vibratome, USA) filled with the same solution and cut in 200 μm transverse slices containing the dorsal hippocampus. Then, the slices were placed in aCSF solution containing (mM): 120 NaCl, 2.8 KCl, 1.25 NaH_2_PO_4_, 26 NaHCO_3_, 20 Glucose, 2 CaCl_2_, 1 MgCl_2_ at 34–35 °C for 45 min. Slices were then left at room temperature until use.

### Whole Cell Patch Clamp Recordings

CA1 pyramidal neurons were visualized with an Olympus BX51WI Microscope (Olympus, Japan) with infrared differential interference contrast (IR-DIC). Neurons were chosen based on the morphology (pyramidal shape) and position in the pyramidal layer. Patch clamp recordings were performed using a Heka EPC10 (HEKA Elektronik, Germany) amplifier with 50 kHz sampling rate and low pass filtered at 3 kHz (Bessel). The slices were placed in the recording chamber filled with aCSF and controlled temperature at 34 °C (Scientifica, UK). Electrodes were fabricated from borosilicate glass (BF150-86-10, Sutter Instruments, Novato CA) with tip resistances of 3–5 MΩ.

Glutamatergic excitatory post-synaptic currents (EPSCs) were evoked with the stimulation of Schaffer collaterals with a concentric bipolar microelectrode (FHC – Bowdoin, ME, USA), connected to a SD9 Grass voltage stimulator (Natus Medical Incorporated, Warwick, RI, USA). We used the minimum voltage necessary to produce the current with maximum amplitude. EPSCs were recorded at holding potentials from − 70 mV to +80 mV, with increments of +30 mV with an internal solution consisting of (mM): 130 CsCl, 10 Hepes, 5 EGTA, 5 phosphocreatine, 4 Mg-ATP, 0.5 Na-GTP, 10 TEA, 5 QX 314 pH adjusted to 7.3 with CsOH and ≈ 290 mOsm/kgH_2_O.

To isolate NMDA currents, we applied the AMPA/kainite antagonist 6,7-dinitroquinoxaline-2,3-dione (DNQX; 10 μM). AMPA/KA currents were obtained by subtracting the currents before and after DNQX, and the remaining currents were mediated by NMDA receptors, confirmed by their sensitivity to the NMDA antagonist DL-AP5. The NMDA-AMPA ratio was obtained in the same cell by dividing the current evoked at +80 by the current evoked at −70mV. Short-term depression was evaluated with a train of five EPSCs delivered at 20 Hz. Glutamatergic currents were recorded in the presence of the GABAA antagonist picrotoxin (20 μM).

Spontaneous GABAergic currents were recorded in the presence of DNQX with an internal solution consisting of (mM): 145 KCl, 10 Hepes, 0.5 EGTA, 10 phosphocreatine, 4 Mg-ATP, 0.3 Na-GTP, adjusted to pH 7.3 with KOH and ≈ 290 mOsm/kgH_2_O. Spontaneous inhibitory postsynaptic currents (sIPSCs) were recorded for 10 minutes. We then applied tetrodotoxin (TTX, 0.5μM) to block action potentials and record miniature IPSCs (mIPSCs).

Series resistance (<20 MΩ) was compensated in 60%. Any neuron with series resistance increased over 20% during experiments were discarded. Voltages were corrected off-line for a liquid junction potential for each internal solution calculated with Clampex software (Molecular Devices).

### Data Analysis

Data was analyzed using Mini Analysis (Synaptosoft 6.0.3, Fort Lee, NJ, USA) and custom written routines in IgorPro (Wavemetrics, Portland, OR, USA) and Matlab (MathWorks, Natick, MA, USA) and GraphPad Prism 6.0 (GraphPad Software, La Jolla CA, USA). The peak of the EPSCs were used to build IV relationships to estimate the slope conductances using the inward part of the relationship for the AMPA/KA current and the outward for the NMDA currents.

Histograms were built with same fixed bins for different groups of cells and fitted with single Gaussian functions. Analysis of decay kinetics for inhibitory currents was performed by Mini Analysis group analysis with individual currents fitted with double exponential functions. Fast and slow time constants were presented as average and their cumulative distributions were compared.

Data are reported as mean ± SEM. Unpaired t-test was used to compare the means, setting the significance level (p) below 0.05. We used two-way ANOVA to compare the train of EPSCs. Outliers were identified by the ROUT method of nonlinear regression (Q = 1%).

### Drugs

The following drugs were used: picrotoxin (Sigma, 20 μM), DNQX (Sigma, 10 μM) and tetrodotoxin citrate (TTX); Alomone Labs, 0.5 μM). DNQX was dissolved in DMSO and then added to bath from fresh stock solutions. The final concentration of DMSO in the experiments was 0.1% and since we did not find differences between DMSO and aCSF, we used only aCSF as control vehicle. All salts were of reagent grade.

## Results

### High-intensity noise does no change glutamatergic excitatory transmission

We compared the Schaffer-collateral AMPA/KA-mediated excitatory post-synaptic currents in both sham and stimulated groups (Figure 1A). We found that AMPA/KA EPSCs from pyramidal neurons from hippocampi of stimulated animals have similar amplitudes (control = −695 ± 71 pA; stimulated = −617 ± 111 pA; p = 0.42; unpaired t-test, N= 15 and 14, respectively) (Fig 1B.) and slope conductances (control = 6.3 ± 0.8nS; stimulated = 8.1 ± 0.9 nS; p = 0.19; N= 15 and 14, respectively). IV relationships of the AMPA/KA currents (DNQX-sensitive) were also similar, showing a small inward rectification in both groups (Figure 1C). On the other hand, we found increased facilitation of the EPSCs during trains of stimulation in synapses from stimulated animals (20 Hz; Pn/P1 F(1,24) = 5.3; p=0.03, Two-Way ANOVA N= 15 and 8, respectively) (Fig. 1D,E).

**Figure 1.**
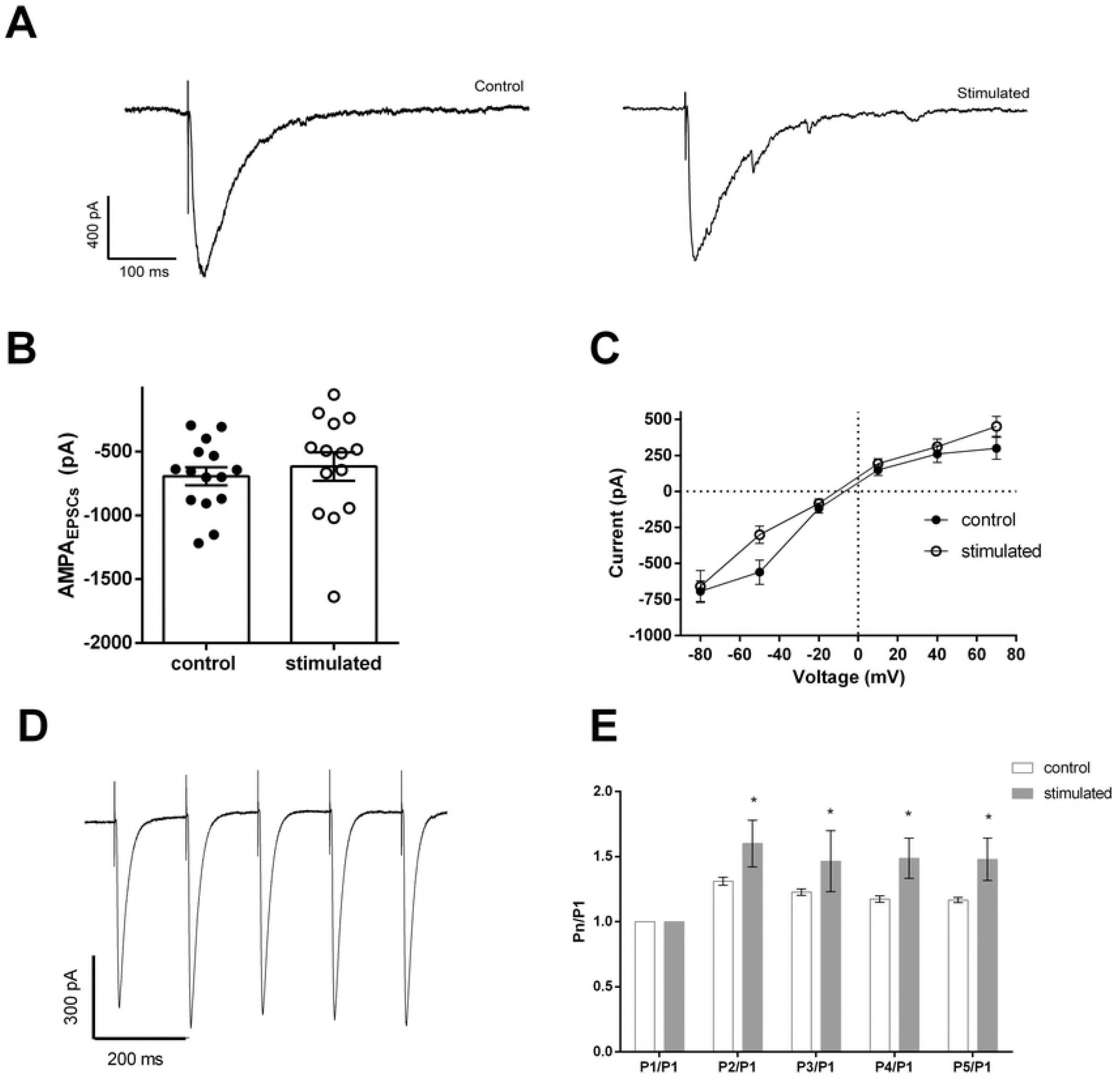
Evoked excitatory AMPA/KA post-synaptic currents (EPSCs) in CA1 pyramidal cells. A. EPSCs recorded in the presence of Picrotoxin (20 μM) at −80 mV from control and stimulated animals. D. Mean AMPA/KA current amplitudes evoked at −80 mV. C. IV relationships for AMPA/KA currents. C. EPSCs in response to a 20 Hz train of stimuli. D. P1/Pn relationships.

Next, we isolated NMDA-mediated currents using DNQX (10 μM) in order to check whether high-intensity sound could affect these currents, which are crucial for LTP induction in the Schaffer-CA1 synapses (Fig. 2A). We found that the amplitudes of NMDA EPSCs were not significantly different in sound stimulated animals (control = 317.5 ± 44.6 pA and stimulated = 261 ± 35.5 pA; p = 0.33; unpaired t-test, N= 17 and 15, respectively. Figure 2B). The IV relationships of the NMDA currents were similar (Figure 2C) and their slope conductances, obtained in the IV were also similar (controls = 4.1 ± 0.8 nS; stimulated = 3.1 ± 0.82 nS; p = 0.42; N= 17 and 15, respectively). Finally, we did not observe significant differences in the NMDA/AMPA ratio between cells from controls and stimulated animals (controls = 0.58 ± 0.09 and stimulated = 0.85 ± 0.3; p = 0.4; unpaired t-test, N= 17 and 15, respectively).

**Figure 2.**
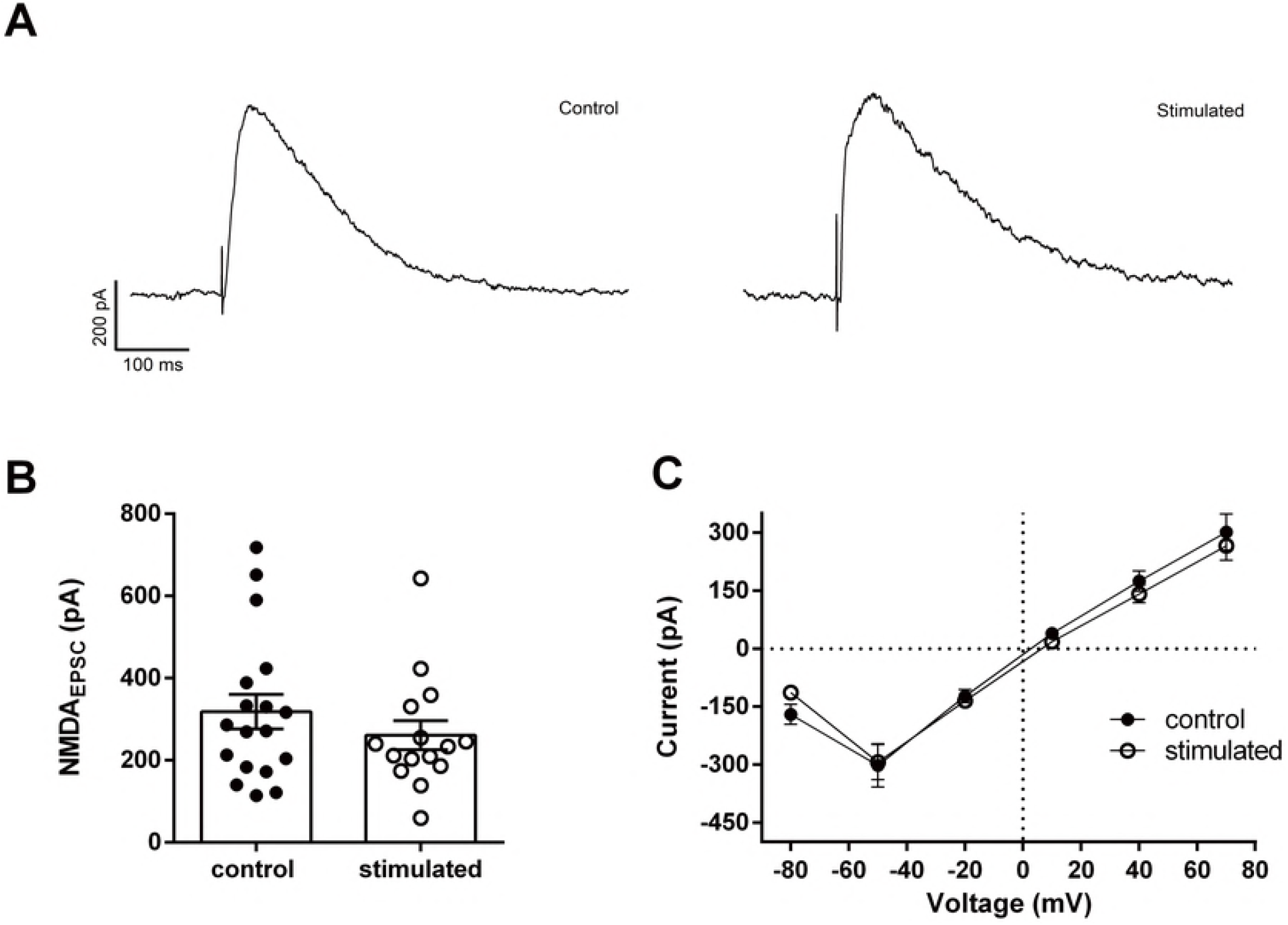
Evoked excitatory NMDA post-synaptic currents (EPSCs) in CA1 pyramidal. A. EPSCs recorded in the presence of Picrotoxin (20 μM) and DNQX (10 μM) at +50 mV from control and stimulated animals. B. Mean NMDA current amplitudes evoked at +50 mV. C. IV relationships for DNQX-sensitive NMDA currents.

### High-intensity noise potentiates inhibitory GABAergic transmission

We then investigated the GABAergic transmission on hippocampal CA1 pyramidal neurons, recording the spontaneous GABaergic IPSCs (sIPSCs) in slices from control and stimulated animals (Figure 3A). The frequency of the sIPSCs was not significantly different between neurons from control and stimulated animals (control: 3.9 ± 0.6 Hz; stimulated: 3.6 ± 0.5 Hz; p=0.7; n= 19 and 26 respectively. Figure 3B) but the mean amplitudes were bigger in neurons from stimulated animals (control: −76 ± 4 pA; stimulated: −98 ± 8 pA; p = 0.05; n= 17 and 25 respectively. Figure 3C*i*). The distribution of the amplitudes followed a bell-shape curve with a skewness toward bigger amplitudes (Figure 3C*ii*). Simple Gaussian functions were fitted to the frequencies histograms, and they produce curves with significant different means (control: −57.6 ± 1.5 pA; stimulated: −72.5 ± 2.5 pA; p<0.0001). On the other hand, the half-width of the sIPSCs was shorter in stimulated animals (control: 3.5 ± 0.3 ms; stimulated: 2.8 ± 0.1 ms; p = 0.026; n= 17 and 25 respectively. Figure 3D).

**Figure 3.**
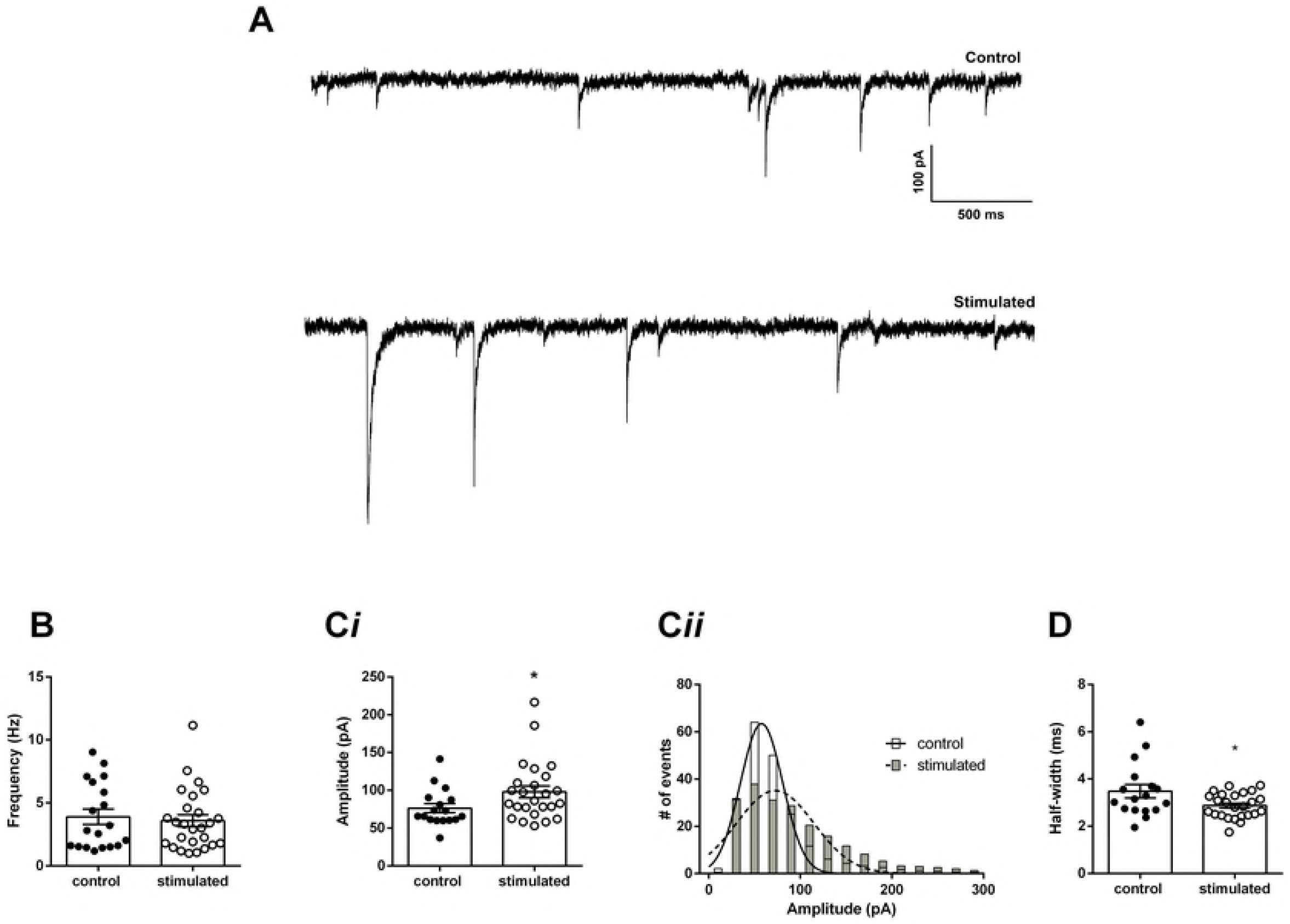
Spontaneous inhibitory post synaptic currents (sIPSCs) in CA1 pyramidal cells. A. sIPSCs recorded in the presence of DNQX (10 μM) at −70 mV, from control and stimulated animals. B. Mean frequency of IPSCs from controls and stimulated animals. Ci. Mean current amplitudes and, Cii, Amplitude histograms of spontaneous currents from control and stimulated animals (error bars were omitted for clarity). D. Mean half-widths of recorded IPSCs from control and stimulated animals. *p<0.05.

sIPSCs are affected by the firing of GABAergic neurons. In order to study the properties of the GABAergic synapses we applied TTX to block action potential firing, and recorded the action-potential independent mIPSCs (Figure 4A). The frequency of mIPSCs was smaller than of the sIPSCs, reflecting the spontaneous action-potential-independent release of GABA. Again, we did not observe differences in the frequency of mISPCs from control and stimulated animals (control: 1.5 ± 0.2 Hz; stimulated: 1.2 ± .1 Hz. P = 0.28; N=22 and 13 respectively. Figure 4B). However, the amplitude of the mIPSCs was significantly bigger in CA1 pyramidal neurons from stimulated animals (control: −75.2 ± 3 pA; stimulated: −98.7 ± 5 pA; P = 0.0003; N=22 and 13 respectively. Figure 4C*i*). The amplitude distribution of the mIPSCs amplitude was fitted with a Gaussian function, and the fits had significantly different means (control: −60.4 ± 1 pA; stimulated: −78.1 ± 1.3 pA; p<0.0001;. Figure 4C*ii*). The half-widths were similar (control: 2.8 ± 0.1 ms; stimulated: 2.8 ± 0.1 ms; P=0.6; N=22 and 13 respectively. Figure 4D). On the other hand, a more detailed analysis of the decay time of the mIPSCs showed that the fast decay time constant of mIPSCs was slightly, but significantly faster in neurons of stimulated animals (control 2.8 ± 0.08 ms; stimulated: 2.4 ± 1.3 ms; P = 0.004; N=22 and 13 respectively. Figure 4E) while the slow decay time constant was not different (control 20.7 ±1.2 ms; stimulated: 22.4 ± 0.8 ms; P = 0.3; N=22 and 13 respectively. Figure 4F). The fast component comprised of 49.6 ± 0.7 % of the total decay time in control animals and 51.9 ± 0.9 % in stimulated animals (P=0.07).

**Figure 4.**
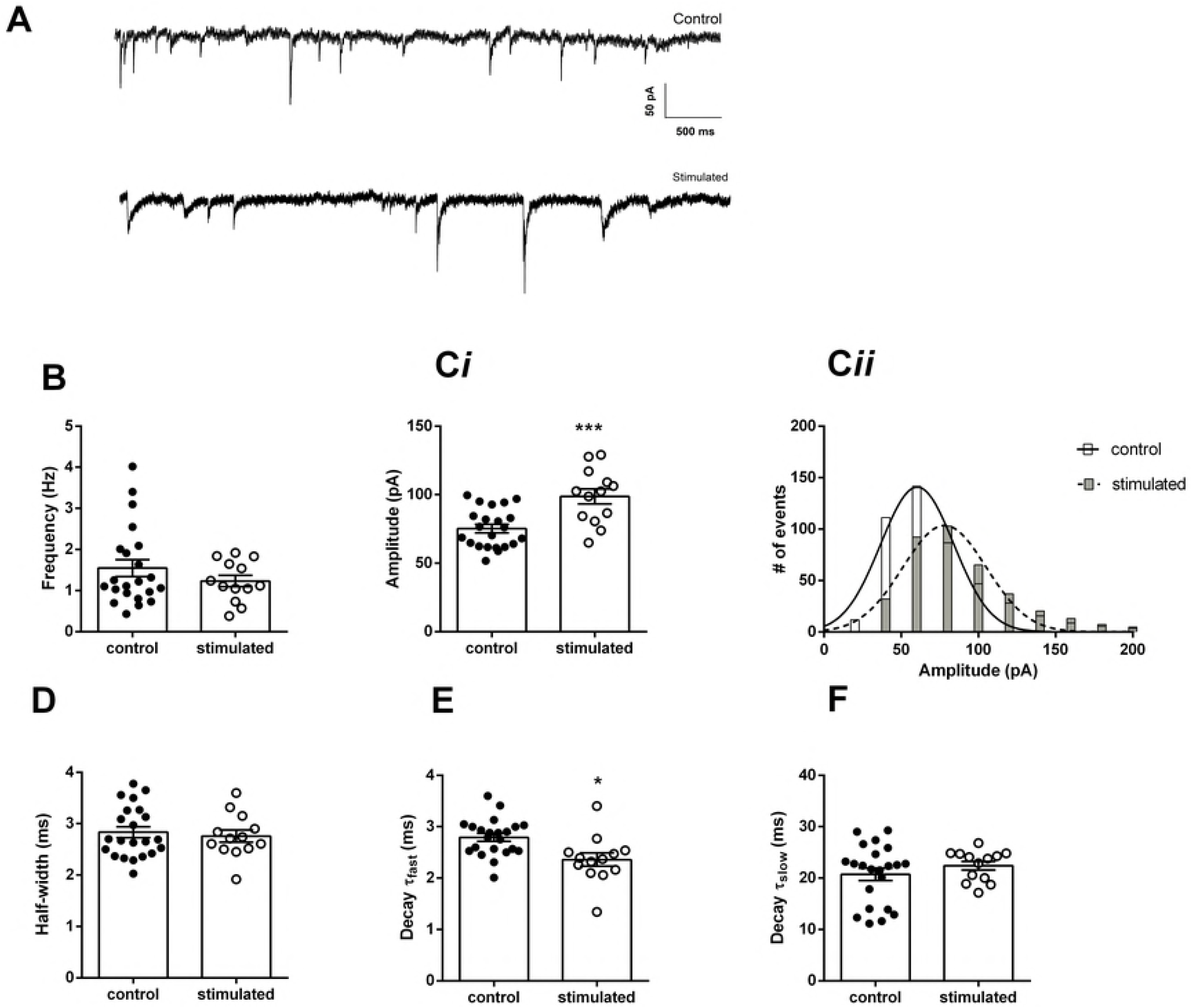
Spontaneous miniature inhibitory post synaptic currents (mIPSCs) in CA1 pyramidal cells. A. Current traces recorded in the presence of DNQX (10 μM) and TTX (1 μM) at −70 mV, from control and stimulated animals.. B. Mean frequency of mIPSCs from controls and stimulated animals. Ci. Mean current amplitudes and Cii, amplitude histograms of spontaneous currents from control and stimulated animals (error bars were omitted for clarity). D. Mean half-widths. E. Mean fast decay time constants. F. Mean slow decay time constants. *p<0.05, ***p<0.001.

## Discussion

Here we showed that GABAergic transmission is potentiated in the CA1 area of the hippocampus of rats after a protocol of 20 episodes of one minute of high-intensity noise, delivered during the period of 10 days. On the other hand, we did not observe any change in the excitatory glutamatergic transmission via both AMPA/kainate and NMDA receptors, except an increase in the short term facilitation during a 20 Hz train. Our findings add to others and our previous findings, showing that the acoustic environment affects the hippocampus.

We have shown previously that the Schaffer-CA1 LTP was inhibited in animals one week after exposure to the same sound exposure protocol used in this investigation [18]. We also observed a diminished h current in CA1 pyramidal neurons, which hyperpolarized the membrane, and increased the membrane time constant [19]. The membrane hyperpolarization could negatively affect action potential firing in response to the train of EPSPs used to induce LTP, affecting the coincidence of pre and post-synaptic firing necessary for inducing associative LTP in the hippocampus [22]. On the other hand, the increased membrane time constant can potentially increase dendritic summation of the EPSPs [23, 24], which could have a facilitatory effect on LTP. However, we have not investigated the effects of long-term sound exposure on the inhibitory and excitatory synapses on CA1 pyramidal neurons. The inhibitory effect on LTP could be caused by a decreased glutamatergic transmission via NMDA receptors, which are necessary for inducing associative LTP in the Schaffer-CA1 pathway [22]. However, we did not observe any change in the NMDA receptor mediated glutamatergic transmission. Decreased neurotransmission trough AMPA/kainate receptors could also decrease the probability of inducing LTP, but we also did not observe any change in this transmission. We even found a bigger facilitation of the EPSCs amplitudes during a 20 Hz train. Thus, we cannot explain the decrease in LTP by long-term sound exposure by a reduction in the glutamatergic transmission.

The hippocampus has several types of GABAergic interneurons, which provide strong inhibition controlling the excitability of the pyramidal neurons by both GABA_A_ and GABA_B_ receptors [25, 26]. In hippocampal pyramidal neurons, GABAergic synapses are both in dendrites along the excitatory terminals, and in the proximal dendrites and cell bodies [27]. Activation of dendritic GABAergic synapses increases the threshold for firing of pyramidal neurons, while the activation of proximal and somatic synapses reduces the maximal firing [26]. We found an increase in the amplitude of GABAergic currents after exposure to our high intensity sound protocol, in both the spontaneous and the action potential-independent miniature IPSCs. The lack of change in the frequency and the persistence of this effect after perfusion of TTX strongly suggests that these effects are post-synaptic, on the GABA_A_ receptor level, and not on the release probability of GABAergic vesicles or interneuron firing. The small change in the decay kinetics of the mIPSCs from stimulated animals suggests a change in receptor composition.

These stronger GABAergic currents could reduce the excitability of the pyramidal neuron during the induction of LTP reducing its firing and impairing LTP induction, which depends upon pre-and post-synaptic firing. Somatic inhibition is more effective in impairing pyramidal neuron firing than dendritic inhibition [26] and most of our recorded spontaneous GABAergic currents are probably of somatic origin, because dendritic currents are largely filtered by the dendritic cable resistance [28]. Interestingly, dendritic GABAergic IPSCs were much more sensitive to TTX than somatic mIPSCs [29] suggesting that our mIPSCs were from somatic origin. Additionally somatic whole cell recording seems to increase the probability of GABA release from somatic synapses [30]. This increased GABAergic neurotransmission could also have a protective effect against the development of audiogenic limbic seizures, since the GABAergic inhibition strongly controls the excitability of the hippocampal circuit and GABAergic inhibition leads to development of hippocampal seizures [31]. Interestingly, GABAergic inhibition on CA1 pyramidal neurons is decreased in the hippocampus of a strain susceptible to audiogenic seizures [32].

This increased GABAergic tone is a potential contributor to the inhibited LTP after high intensity sound stimulation, however, at this moment we cannot make any causal conclusion about the mechanism of LTP inhibition. It is possible that this effect is a compensatory mechanism for the increased firing of the pyramidal neurons after high-intensity sound stimulation, for instance [19]. More studies are being carried out in order to elucidate the mechanism of LTP inhibition by high intensity sound exposure. Nevertheless, our findings show that the hippocampal inhibitory neurotransmission is affected by high intensity sound exposure, which can have relevant consequences to the hippocampal function in animals and humans subjected to constant or even episodic loud sound exposure in their environments.

## Acknowledgements

We thank the technical assistance of Mr. J. Fernando Aguiar.

## References

1. Ising H, Kruppa B. Health effects caused by noise: Evidence in the literature from the past 25 years. Noise Health. 2004; 6:5–13

2. Hoffer ME Levin BE, Snapp H, Buskirk J, Balaban C. Acute findings in an acquired neurosensory dysfunction. Otol Neurotol Neurosci. 2018; Epub ahead of print. doi: 10.1002/lio2.231.

3. Bhatt JM, Lin HW, Bhattacharyya N. Prevalence, Severity, Exposures, and Treatment Patterns of Tinnitus in the United States. JAMA Otolaryngol Head Neck Surg. 2016; 21

4. Coelho CB, Sanchez TG, Tyler RS. Hyperacusis, sound annoyance, and loudness hypersensitivity in children. Prog Brain Res. 2007; 166:169–78

5. Helfer TM, Jordan NN, Lee RB, Pietrusiak P, Cave K, Schairer K.. Noise-induced hearing injury and comorbidities among postdeployment U.S. Army soldiers: April 2003-June 2009. Am J Audiol. 20:33–41 (2011).

6. Lercher, P., Evans, G.W., Meis, M. Ambient noise and cognitive processes among primary schoolchildren. Env Behav. 2003; 35:725–735

7. Zocoli, A.M., Morata, T.C., Marques, J.M. & Corteletti, L.J. Brazilian young adults and noise: attitudes, habits, and audiological characteristics. Int. J. Audiol. 2009; 48:692–9

8. Kraus KS, Canlon B. Neuronal connectivity and interactions between the auditory and limbic systems. Effects of noise and tinnitus. Hear Res. 2012; 288:34–46

9. Zhao H, Wang L, Chen L, Zhang J, Sun W, Salvi RJ, Huang YN, Wang M, Chen L. Temporary conductive hearing loss in early life impairs spatial memory of rats in adulthood. Brain Behav. 2018; 8(7):e01004. doi: 10.1002/brb3.1004.

10. Squire LR, Schmolck H, Stark SM. Impaired auditory recognition memory in amnesic patients with medial temporal lobe lesions. Learn Mem. 2001; 8:252–6

11. Tamura R, Ono T, Fukuda M, Nakamura K. Recognition of egocentric and allocentric visual and auditory space by neurons in the hippocampus of monkeys. Neurosci. Lett. 1990; 109:293–298

12. Aronov D, Nevers R, Tank DW. Mapping of a non-spatial dimension by the hippocampal-entorhinal circuit. Nature. 2017; 543:719–722

13. Wang N, Gan X, Liu Y, Xiao Z. Balanced Noise-Evoked Excitation and Inhibition in Awake Mice CA3. Front Physiol. 2017; 8:931. doi: 10.3389/fphys.2017.00931.

14. Cheng L, Wang SH, Chen QC, Liao XM.. Moderate noise induced cognition impairment of mice and its underlying mechanisms. Physiol Behav. 2011; 104:981–8

15. Cheng L, Wang SH, Huang Y, Liao XM. The hippocampus may be more susceptible to environmental noise than the auditory cortex. Hear Res. 2016; 333:93–7.

16. Goble TJ, Møller AR, Thompson LT. Acute high-intensity sound exposure alters responses of place cells in hippocampus. Hear Res. 2009; 253:52–9.

17. Kapolowicz MR, Thompson LT. Acute high-intensity noise induces rapid Arc protein expression but fails to rapidly change GAD expression in amygdala and hippocampus of rats: Effects of treatment with D-cycloserine. Hear Res. 2016; 342:69–79.

18. Cunha AO, de Oliveira JA, Almeida SS, Garcia-Cairasco N, Leão RM. Inhibition of long-term potentiation in the schaffer-CA1 pathway by repetitive high-intensity sound stimulation. Neuroscience. 2015; 310:114–27.

19. Cunha AOS, Ceballos CC, de Deus JL, Leão RM. Long-term high-intensity sound stimulation inhibits h current (I_h_) in CA1 pyramidal neurons. Eur J Neurosci. 2018; 7:1401–1413.

20. de Deus JL, Cunha AOS, Terzian AL, Resstel LB, Elias LLK, Antunes-Rodrigues J, Almeida SS, Leão RM. A single episode of high intensity sound inhibits long-term potentiation in the hippocampus of rats. Sci Rep. 2017; 7:14094. doi: 10.1038/s41598-017-14624-1.

21. Romcy-Pereira, RN Garcia-Cairasco, N. Hippocampal cell proliferation and epileptogenesis after audiogenic kindling are not accompanied by mossy fiber sprouting or Fluoro-Jade staining. Neuroscience. 2003; 119: 533–546.

22. Luscher C, Malenka RC. NMDA Receptor-Dependent Long-Term Potentiation and Long-Term Depression (LTP/LTD). Cold Spring Harb Perspect Biol. 2012; doi:10.1101/cshperspect.a005587.

23. Magee, J. C. Dendritic hyperpolarization-activated currents modify the integrative properties of hippocampal CA1 pyramidal neurons. J. Neurosci. 1998; 18, 7613–7624.

24. Maroso, M., Szabo, G.G., Kim, H.K., Alexander, A., Bui, A. D., Lee, S. H., Lutz, B., & Soltesz, I. Cannabinoid Control of Learning and Memory through HCN Channels. Neuron. 2016; 89:1059–73.

25. Pelkey KA, Chittajallu R, Craig MT, Tricoire L, Wester JC, McBain CJ. Hippocampal GABAergic Inhibitory Interneurons. Physiol Rev. 2017; 97:1619–1747.

26. Pouille F, Watkinson O, Scanziani M, Trevelyan AJ. The contribution of synaptic location to inhibitory gain control in pyramidal cells. Physiol Rep. 2013; 1:e00067. doi: 10.1002/phy2.67.

27. Megías M, Emri Z, Freund TF, Gulyás AI. Total number and distribution of inhibitory and excitatory synapses on hippocampal CA1 pyramidal cells. Neuroscience.;102:527–40. 2001.

28. Williams SR, Mitchell SJ. Direct measurement of somatic voltage clamp errors in central neurons. Nat. Neurosci. 2008; 11:790–798.

29. Cossart R, Hirsch JC, Cannon RC, Dinoncourt C, Wheal HV, Ben-Ari Y, Esclapez M, Bernard C. Distribution of spontaneous currents along the somato-dendritic axis of rat hippocampal CA1 pyramidal neurons. Neuroscience. 2000; 99:593–603.

30. Andrásfalvy BK, Mody I. Differences between the scaling of miniature IPSCs and EPSCs recorded in the dendrites of CA1 mouse pyramidal neurons. J Physiol. 2006.576:191–196.

31. Khazipov R, Khalilov I, Tyzio R, Morozova E, Ben-Ari Y, Holmes GL. Developmental changes in GABAergic actions and seizure susceptibility in the rat hippocampus. Eur J Neurosci. 2004;19:590–600.

32. Cunha AOS, Ceballos CC, de Deus JL, Pena RFO, de Oliveira JAC, Roque AC, Garcia-Cairasco N, Leão RM. Intrinsic and synaptic properties of hippocampal CA1 pyramidal neurons of the Wistar Audiogenic Rat (WAR) strain, a genetic model of epilepsy. Sci Rep. 8:10412. 2018; doi: 10.1038/s41598-018-28725-y..

